# Physioxia Reprograms Glioblastoma Cells Enhancing Migration and Altering Therapeutic Sensitivity

**DOI:** 10.64898/2026.07.05.736632

**Authors:** Natasha Hockaden, Elise O’Herron, Dylan Zhou, Margaret Heffernan, Scott Cooper, Angela Richardson

## Abstract

**Background/Objectives:** Glioblastoma is an aggressive primary brain tumor that develops within a chronically low-oxygen microenvironment, yet most preclinical studies are performed under atmospheric oxygen conditions that poorly reflect in vivo physiology. This study investigated how sustained culture under physiological oxygen tension (physioxia; 5% O□) influences glioblastoma cell behavior, signaling, and therapeutic response.

**Methods:** Multiple patient-derived glioblastoma models were cultured under normoxia (21% O□) or sustained physioxia (5% O□) for at least seven days before experimentation. Cell migration, proliferation, cell cycle distribution, expression of the epithelial-to-mesenchymal transition-associated transcription factor Slug (SNAI2), PDGFRβ-associated signaling, and sensitivity to 5-fluorouracil were evaluated using transwell migration assays, cell counting, flow cytometry, RT-qPCR, immunoblotting, and BrdU incorporation assays. Additional patient-derived cultures established and maintained continuously under physioxia were used to examine the effects of oxygen history.

**Results:** Sustained physioxia consistently increased migration across all glioblastoma models while reducing proliferation in normoxia-adapted cell lines through increased G0/G1 cell cycle arrest. Physioxia significantly increased Slug expression in all models and enhanced PDGFRβ, AKT, and ERK phosphorylation in a cell line-dependent manner. Therapeutic sensitivity to 5-fluorouracil was also altered, with physioxia conferring increased resistance in selected glioblastoma models but not universally. Patient-derived cultures maintained continuously under physioxia retained enhanced migratory capacity and exhibited increased proliferation compared with normoxia, indicating that prior oxygen exposure influences proliferative responses while the pro-migratory phenotype remains conserved.

**Conclusions:** Physiological oxygen tension is a major regulator of glioblastoma cell behavior, influencing migration, proliferation, signaling, and therapeutic response. These findings demonstrate that conventional normoxic culture conditions can obscure biologically relevant phenotypes and support incorporating physioxia into experimental design to improve the physiological and translational relevance of preclinical glioblastoma research.

## Introduction

Glioblastoma (GBM) is the most aggressive primary malignancy of the central nervous system and is characterized by diffuse invasion, extensive cellular heterogeneity, and resistance to therapy [1]. Despite maximal surgical resection followed by radiotherapy and temozolomide, median survival remains approximately 15 months [1, 2]. These poor clinical outcomes highlight the need to better understand the biological mechanisms that regulate GBM progression within the tumor microenvironment. A defining feature of GBM is the presence of heterogeneous oxygen gradients that influence tumor cell signaling, proliferation, invasion, and therapeutic response [3, 4].

Standard cell culture conditions expose cells to atmospheric oxygen (∼21% O□), which substantially exceeds oxygen levels present in normal brain tissue and glioblastoma tumors [5, 6]. Physiological oxygen tension in the brain typically ranges from 2-8% O□, whereas GBM tumors frequently contain regions with oxygen levels approaching 1% due to abnormal vasculature and rapid tumor growth [4]. Consequently, culture under physioxia (∼5% O□) more closely reflects the oxygen conditions experienced by GBM cells in vivo than conventional ambient air culture.

Most studies examining oxygen-dependent tumor biology have focused on acute hypoxia models lasting from minutes to 48 hours [7]. Although these systems have defined important hypoxia-responsive pathways, they primarily capture transient stress responses rather than the stable adaptations associated with chronic low-oxygen exposure in tumors [4–7]. In vivo, glioblastoma cells are subjected to prolonged oxygen limitation that alters proliferation, migration, cell cycle progression, and therapy resistance [4, 5, 8]. Long-term culture under physioxia is therefore required to model the sustained oxygen adaptation that occurs in situ rather than acute hypoxic stress. Prior studies assessing the effect of oxygen tension on GBM have identified alteration in many cellular pathways but even those studies that examined “chronic hypoxia” typically limited the exposure to 48-72 hours which was limited to the incubation time, while cell splitting, passaging, and experiments were typically performed in ambient air [references].

Work by Broxmeyer and colleagues demonstrated that physioxia induces durable changes in cellular behavior, including altered stemness, signaling, and cell cycle regulation [7–10]. These studies further showed that even brief exposure to atmospheric oxygen can rapidly alter cellular phenotypes [8, 10]. Such observations are particularly relevant to GBM, which contains stem-like tumor populations associated with invasion, recurrence, and therapeutic resistance. In neural and endothelial cell systems, prolonged physioxic culture alters differentiation and stress responses, suggesting that conventional normoxic conditions may obscure physiologically relevant tumor behavior [11–14].

Among the signaling pathways influenced by oxygen tension, p53 and PDGFRβ-associated signaling are of particular interest in GBM. The tumor suppressor p53 regulates cellular responses to stress, including DNA damage and hypoxia [15, 16]. TP53 mutations are common in glioblastoma and influence tumor progression, signaling responses, and therapeutic sensitivity [17]. Oxygen tension can modulate p53 activity in a context-dependent manner, with distinct effects observed in wild-type and mutant p53 backgrounds [17, 18].

Platelet-derived growth factor receptor beta (PDGFRβ) and its downstream PI3K/AKT and MAPK/ERK signaling pathways are major regulators of glioblastoma proliferation, survival, cytoskeletal remodeling, and migration [11, 19, 20]. These pathways are sensitive to oxygen availability and redox state [11]. In other tumor systems, physioxia enhances PDGFRβ-dependent signaling and promotes migratory behavior through sustained AKT and ERK activation [11]. Loss or mutation of p53 can further amplify these signaling responses by reducing inhibitory regulation of AKT and ERK pathways under low-oxygen conditions [21, 22].

Together, these studies identify oxygen tension as a major determinant of tumor cell behavior rather than a passive culture variable [7, 8, 11]. Modeling GBM under chronic physioxia may therefore provide a more accurate representation of in vivo signaling states, invasive behavior, and therapeutic response. In this study, we investigated how sustained physioxia (minimum of 7 days, with cells passaged and maintained at physiologic oxygen levels) influences glioblastoma migration, proliferation, cell cycle progression, and signaling compared with ambient air conditions. We further examined the relationship between oxygen tension, p53 status, and PDGFRβ-associated AKT and ERK signaling to define mechanisms that regulate glioblastoma adaptation to the brain microenvironment.

## Methods

### Cell Culture and Materials

GBM43 (RRID:CVCL_E5GD) and GBM10 cells were obtained from the Mayo Clinic. GBM001 cells were a generous gift from Dr. Michael Ivan from the University of Miami. For normoxia experiments cells were cultured in 21% O□. For physioxia experiments, cells were maintained at 5% O for at least 7 days prior to being utilized for any experiments. AR022 and AR023 cells were obtained from patients undergoing surgical resection with informed consent for research use (IRB-04, 25737). Tumors were transported in a hypoxia chamber. Cells were isolated and maintained in either ambient air conditions containing 21% O (normoxia, using a typical biosafety cabinet) or physiological oxygen tension (physioxia, Biospherix Chamber set to 5% O□. For cell maintenance and all experiments, plasticware, media, PBS, drug solutions were acclimated for at least 24 hours to the appropriate oxygen tensions. Once cells were fixed, downstream experiments were carried out in ambient air.

GB43, GBM001, and GB10 were cultured in DMEM/F12 (Cytiva SH30023.01) supplemented with 10% FBS (Corning 35015CV), 1% penicillin-streptomycin (Cytiva SV30079.01), 1% Sodium Pyruvate (Gibco 11360070), and 1% MEM Non-Essential Amino Acids (GIBCO 11140050) at 37°C in 5% CO□.

AR022 and AR023 cells were cultured in DMEM/F12 (Cytiva SH30023.01) supplemented with 2.5% FBS (Corning 35015CV), 1% penicillin-streptomycin (Cytiva SV30079.01), 1% Sodium Pyruvate (Gibco 11360070), and 1% MEM Non-Essential Amino Acids (GIBCO 11140050), B-27™ Supplement minus vitamin A (Invitrogen 12587010), N2™ Supplement minus vitamin A (Invitrogen, 17502048), rhEGF, 20ng/ml (R&D Systems, 236-EG), rhFGF, 20ng/ml (R&D Systems, BT-FGFHS) at 37°C in 5% O□.

### Proliferation

Cells were maintained in standard tissue culture flasks and passed at 80% confluence. Media was refreshed every 3 days to maintain optimal growth conditions. At each time point, cells were harvested using trypsinization and resuspended in fresh media. Cell viability and total counts were determined using a hemocytometer and trypan blue dye.

### Chemotherapy response

Cells were seeded into 96 well plates at densities optimized for each cell line (GBM43: 3,000 cells/well; GBM10: 3,000 cells/well; GBM001: 5,000 cells/well; AR022: 2,000 cells/well; AR023: 2,000 cells/well) and allowed to adhere overnight in either physioxia or normoxia. Cells were then treated with 5-fluorouracil (5-FU) (Thermo Scientific A13456.06) at the indicated concentrations, diluted in standard culture medium specific to each cell line. Following 48hr of drug exposure under standard culture conditions and the appropriate oxygen tension, cell proliferation was assessed using a BrdU ELISA kit (Abcam, ab126556) according to the manufacturer’s protocol. BrdU labeling times were optimized for each cell line (4hr for GBM43 and GBM10; 20hr for GBM001, AR022, and AR023). Absorbance was measured at 450nm, with each well read in triplicate. Background signal from no-cell control wells was subtracted, and BrdU incorporation was normalized to the mean absorbance of vehicle-treated controls at the same oxygen tension.

### Cell Cycle

Cells were cultured under standard conditions at the appropriate oxygen tension for each cell line. GBM43, GBM10, and GBM001 cells were seeded in 10-cm tissue culture dishes at a density of 300,000 cells per dish. AR022 and AR023 cells were seeded in T180 flasks at a density of 600,000 cells per flask. Cells were harvested at time points corresponding to the divergence of normoxic and physioxic growth curves for each cell line (day 3 for GBM43; day 7 for GBM10 and GBM001). Cells were fixed in 10% neutral buffered formalin and stained with FxCycle™ Far Red Stain (Invitrogen, F10348) according to the manufacturer’s protocol. Flow cytometry analysis was performed using 633/5 nm excitation and emission. Cell cycle distribution was determined by DNA content, and the G0/G1, S, and G2/M phases were identified by histogram analysis.

### RT- qPCR

Total RNA was isolated using the RNeasy® Mini Kit (Qiagen 74106). RNA was quantified by nanodrop and reverse transcribed using the High-Capacity RNA-to-cDNA Kit (Applied Biosystems 4387406). Quantitative PCR was performed using a SYBR green master mix (Applied Biosystems A25742) along with gene specific primers using QuantStudio6 (Applied Biosystems, RRID:SCR_020239). Primers used included GAPDH forward 5’- TCAAGGCTGAGAACGGGAAG -3’ and reverse 5’-CGCCCCACTTGATTTTGGAG -3’ and Slug forward 5’- TCGGACCCACACATTACCTT-3’ and reverse 5’- TGACCTGTCTGCAAATGCTC-3’.

### Immunoblotting

Immunoblotting was performed using standard sodium dodecyl sulfate polyacrylamide gel electrophoresis techniques with RIPA buffer (50mM Tris, 150mM NaCl, 1mM EDTA, 1% NP-40, 1% sodium deoxycholate, 1% SDS) used to lyse cells as previously described [23]. Antibodies used for immunoblotting included Histone 3 (Cell Signaling Technology 9715, RRID: AB_331563), pPDGFRβ (Y751) (Cell Signaling Technology 4549, RRID:AB_1147704), PDGFRβ (Cell Signaling Technology 3162, RRID:AB_331111), pAKT (S473) (Cell Signaling Technology 4060, RRID:AB_2315049), AKT (Cell Signaling Technology 4691, RRID:AB_915783), pERK (T202/Y204) (Cell Signaling Technology 9101, RRID:AB_331646), ERK (Cell Signaling Technology 9102, RRID:AB_330744), P53, pP53 (S15), β-catenin, and pβ-catenin (S675).

### Transwell Migration

Cells were deprived of FBS for 18 hours before performing a transwell migration assay using ThinCert Boyden chamber (Greiner One 662638). After starvation, conducted at the appropriate oxygen tension, 50,000 cells were plated into each insert in serum-free medium in a 24-well plate. Complete growth medium (with 10% FBS for GBM001, GBM10, and GBM43 cells and 2.5% FBS for AR022 and AR023 cells) was placed in the lower chamber to serve as a chemoattractant. Cells were plated in triplicate for each condition and were allowed to migrate for 24 hrs where they were then stained and analyzed as previously described [24].

### Statistical Analysis

All quantitative data are presented as mean ± standard error of the mean (SEM), unless otherwise indicated. Statistical analyses were performed using GraphPad Prism (version 10.4.2, San Diego, CA). The number of biological replicates (independent experiments or patient-derived cell lines) and technical replicates for each assay are specified in the corresponding figure legends. For comparisons between two groups (e.g., normoxia vs physioxia), unpaired two-tailed Student’s t tests were used when data were normally distributed, and variances were comparable. For experiments involving more than two groups or multiple conditions (e.g., dose-response analyses, time-course proliferation assays, or comparisons across multiple cell lines), one-way or two-way analysis of variance (ANOVA) was performed as appropriate. Two-way ANOVA was used for experiments assessing the effects of oxygen tension and an additional variable (e.g., drug concentration or time). When ANOVA revealed a significant main effect or interaction, post hoc multiple-comparison testing was conducted using Tukey’s or Sidak’s correction, as specified in the figure legends.

Transwell migration assays, proliferation assays, cell cycle distributions, gene expression analyses, and immunoblot quantifications were analyzed using biological replicates as the unit of analysis. For flow cytometry-based cell cycle analysis, percentages of cells in each phase (G0/G1, S, and G2/M) were compared between oxygen conditions using unpaired two-tailed t tests or ANOVA, depending on the number of groups analyzed.

For chemotherapeutic response assays, dose-response curves were generated, and percent survival was calculated relative to untreated controls for each oxygen condition. Statistical comparisons between normoxia and physioxia at individual drug concentrations were performed using two-way ANOVA with multiple-comparison correction. No curve fitting-based IC comparisons were performed unless explicitly stated.

RT-qPCR data were analyzed using the ΔΔCt method, with expression levels normalized to housekeeping genes and presented relative to normoxic controls. Log-transformed ΔCt values were used for statistical testing to satisfy normality assumptions.

Western blot densitometry was performed using ImageJ or equivalent software. Phosphorylated protein levels were normalized to total protein or loading controls as indicated. Normalized values were analyzed using unpaired t tests or ANOVA as appropriate.

All statistical tests were two-sided, and p-values < 0.05 were considered statistically significant. Exact p-values and statistical tests used are reported in the figure legends. No statistical methods were used to predetermine sample size, but sample sizes are consistent with those commonly reported in the field. No data was excluded from analysis unless pre-specified experimental quality control criteria were not met.

## Results

### Figure 1: Physioxia enhances transwell migration of glioblastoma cells

Physioxia has been increasingly recognized as an important regulator of tumor cell behavior [8, 11]. Prior studies in multiple cancer types, including breast, prostate, colorectal, and pancreatic cancers, have demonstrated that culturing cells under physioxia (typically 3-5% O□) enhances migratory and invasive capacity compared with ambient air conditions [11, 14, 25]. These effects have been attributed to oxygen-sensitive signaling pathways that regulate cytoskeletal dynamics, extracellular matrix interactions, and motility-associated gene expression [26]. Acute hypoxia has been associated with increased glioblastoma cell migration and invasion through activation of hypoxia-inducible signaling pathways in GBM [4]; however, the impact of sustained physioxia on the migratory behavior of patient-derived glioblastoma cells remains incompletely characterized. To determine the effect of physioxia oxygen tension on glioblastoma cell motility, transwell migration assays were performed using GBM001, GBM10, and GBM43 cells cultured under ambient air/normoxia (∼21% O□) or physioxia (5% O□). Across all three glioblastoma lines, exposure to physioxia resulted in a significant increase in migratory capacity compared with cells maintained under ambient oxygen conditions (Figure 1).

**Figure 1:**
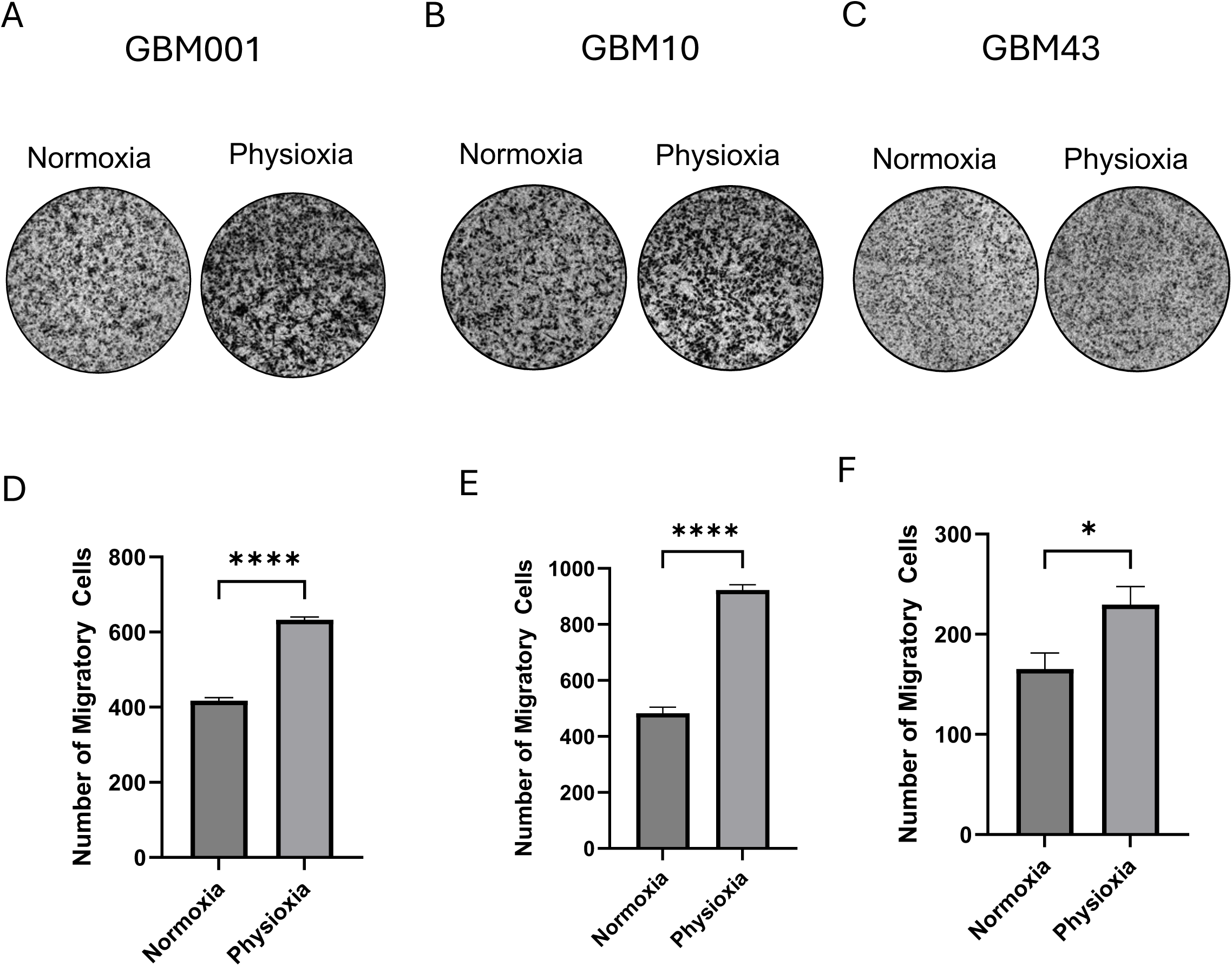
Physioxia enhances glioblastoma cell migration. (A-C) Representative images of migrated (A) GBM001, (B) GBM10, and (C) GBM43 cells following a 24-h Transwell (Boyden chamber) migration assay under normoxic (21% O□) or physioxic conditions (5% O□). (D-F) Quantification of migrated cells demonstrating significantly increased migration under physioxia compared with normoxia in (D) GBM001, (E) GBM10, and (F) GBM43 cells. Data are presented as mean ± SEM from three independent experiments (n = 3); individual experiments were performed in triplicate. P-values calculated by one-way ANOVA where * p< 0.05 and **** p<0.0001

Quantitative analysis revealed that GBM001 cells exhibited a robust elevation in transwell migration under physioxia relative to ambient air (Figure 1D). Similarly, GBM10 cells demonstrated a marked increase in the number of migrated cells when cultured at 5% oxygen Figure 1E). GBM43 cells also showed a significant enhancement in migration under physioxic conditions, indicating that the pro-migratory effect of physioxia is conserved across genetically distinct glioblastoma models (Figure 1F).

Together, these findings demonstrate that physioxia significantly promotes glioblastoma cell migration, extending prior observations from other cancer types to patient-derived GBM models and highlighting oxygen tension as a key microenvironmental determinant of glioblastoma invasiveness. Given the pronounced increase in migratory behavior observed under physioxia, we next examined whether physioxia also affects glioblastoma cell growth characteristics. The following section investigates the effects of physioxia versus ambient air on cell proliferation and cell cycle distribution to determine whether enhanced migration is associated with concurrent changes in proliferative dynamics.

### Figure 2: Physioxia reduces glioblastoma cell proliferation over extended culture

Physioxia has been reported to modulate tumor cell proliferation in a context-dependent manner [27]. Previous studies in multiple cancer types, including breast cancer and other solid tumors, have shown that culturing cells under physioxia can suppress proliferation while promoting invasive or stem-like phenotypes, suggesting a potential trade-off between migratory capacity and cell cycle progression [11, 25]. These observations provided the rationale for examining whether the increased migration observed under physioxia in glioblastoma cells was accompanied by altered proliferative behavior.

**Figure 2:**
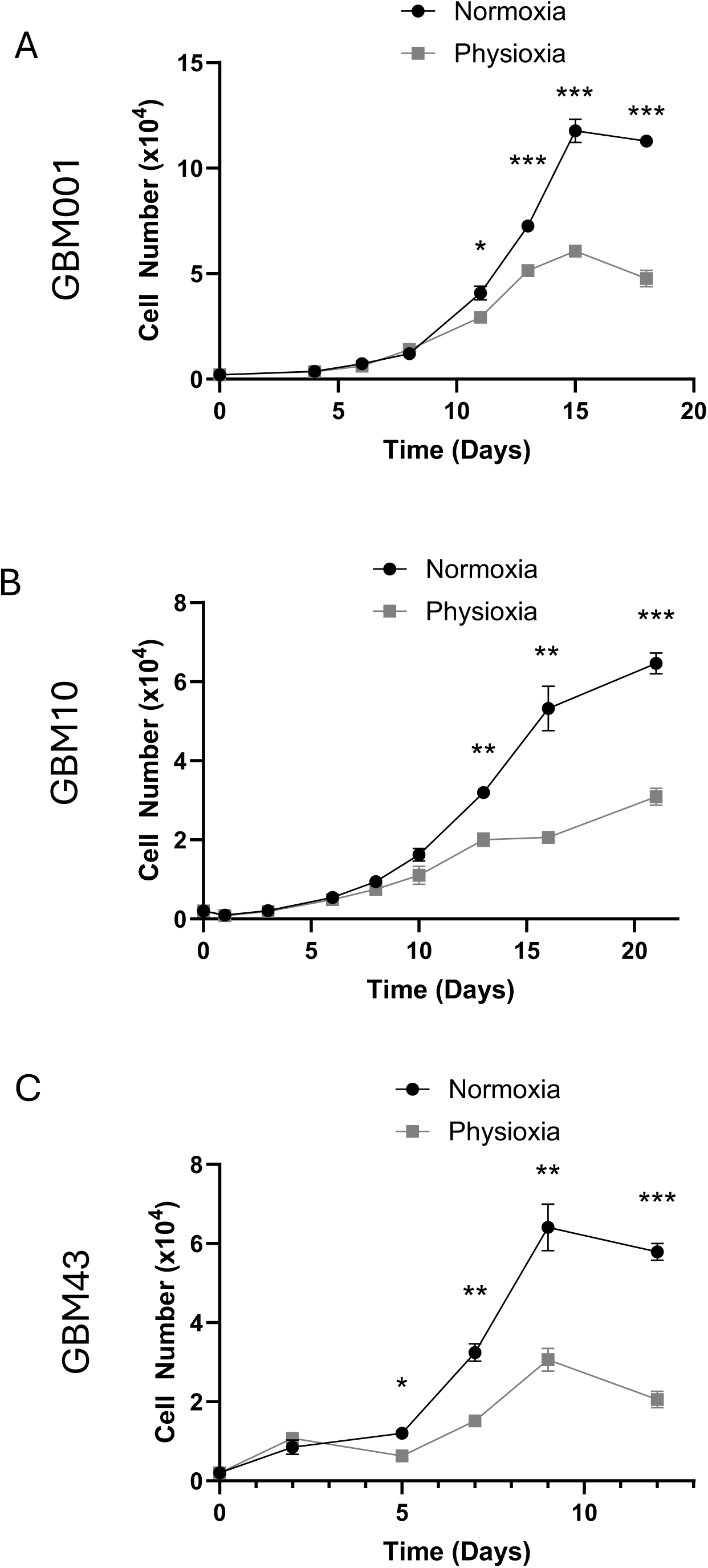
Physioxia reduces long-term proliferation of glioblastoma cells. (A-C) Proliferation of (A) GBM001, (B) GBM10, and (C) GBM43 cells was assessed by cell counting over a 22-day period under normoxic or physioxic conditions. All three cell lines exhibited decreased proliferation when cultured in physioxia compared with normoxia. Data are presented as mean ± SEM from three independent experiments (n = 3). P-values calculated by one-way ANOVA where * p< 0.05, ** p<0.01, *** p<0.001

To assess the effect of oxygen tension on long-term cell growth, GBM001, GBM10, and GBM43 cells were cultured under ambient air or physioxia, and total cell numbers were quantified over a 12-21 day period. Across all three glioblastoma lines, physioxia resulted in a significant reduction in cell accumulation compared with ambient air, although the onset and magnitude of this effect varied by cell line (Figure 2).

In GBM001 and GBM10 cells, a decrease in proliferation under physioxia became evident by Day 7 and persisted throughout the remainder of the culture period (Figure 2A-B). In contrast, GBM43 cells exhibited an earlier sensitivity to physiological oxygen tension, with significantly reduced cell numbers observed as early as Day 3 under physioxic conditions (Figure 2C). These findings indicate that while physioxia consistently suppresses glioblastoma cell proliferation, the temporal response to reduced oxygen tension is cell line-dependent. These findings also demonstrate the need for studies in sustained low oxygen tension to elucidate the impact of oxygen tension on phenotypic readouts.

Given the observed reduction in cell proliferation under physioxia, we next sought to determine whether these effects were associated with alterations in cell cycle progression. Based on prior reports linking physioxia to cell cycle regulation and quiescence, flow cytometry-based cell cycle analysis was performed at the time points corresponding to the onset of proliferation differences in each cell line to further characterize the mechanisms underlying physioxia-induced growth suppression.

### Figure 3: Physioxia induces cell cycle arrest in glioblastoma cells

In GBM001 cells analyzed at Day 7, physioxia resulted in a significant increase in the proportion of cells in the G0/G1 phase, accompanied by a corresponding decrease in S-phase cells compared with ambient air conditions (Figure 3A). A similar pattern was observed in GBM10 cells at Day 7, where culture under physioxia led to an accumulation of cells in G0/G1 and a reduction in S-phase entry, consistent with impaired cell cycle progression (Figure 3B).

**Figure 3:**
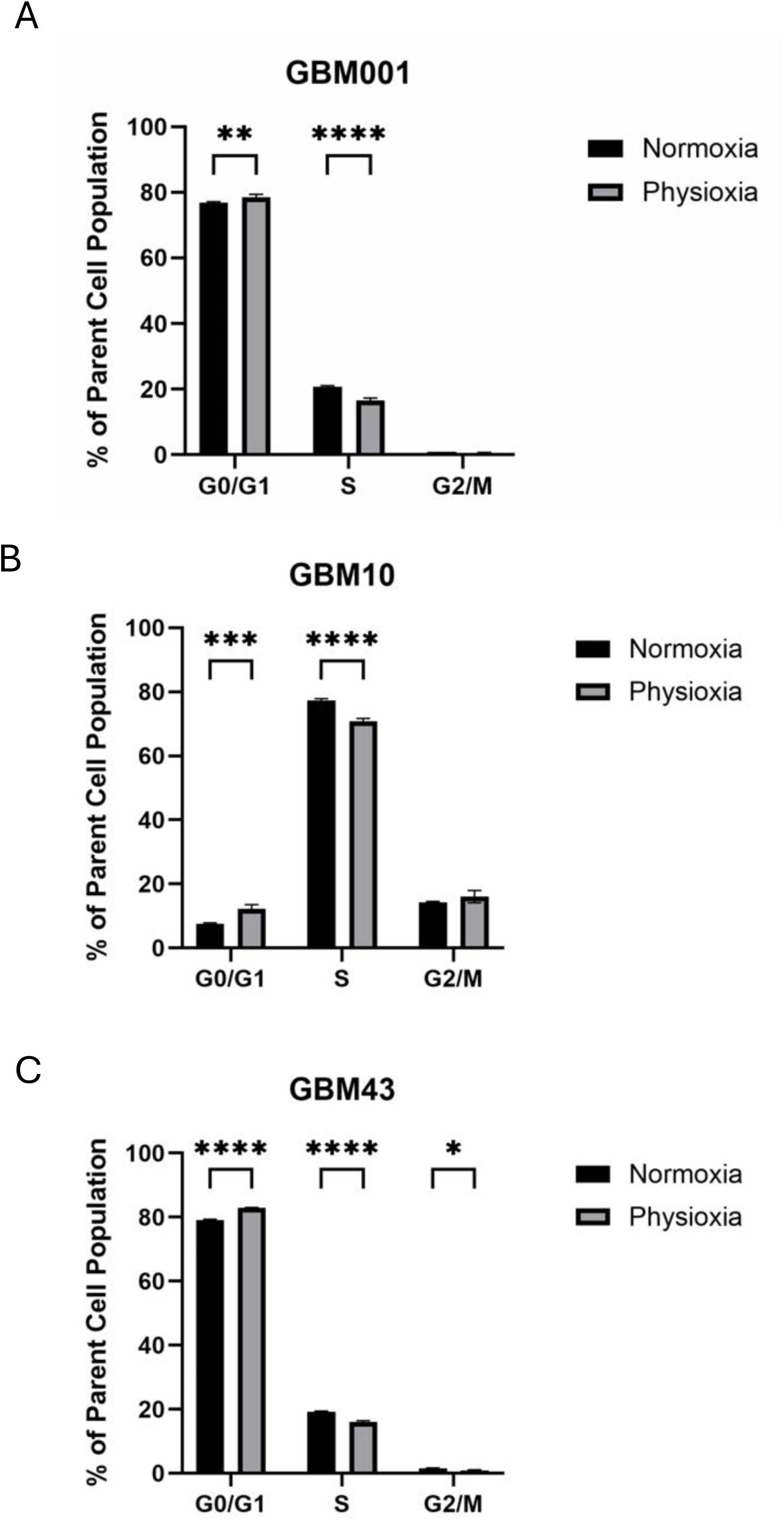
Physioxia alters cell cycle distribution in glioblastoma cells. (A-C) Cell cycle distribution of (A) GBM001 (Day 7), (B) GBM10 (Day 7), and (C) GBM43 (Day 3) cells was analyzed by flow cytometry under normoxic and physioxic conditions. Physioxia resulted in an increased proportion of cells in the G0/G1 phase with a corresponding reduction in S phase in GBM001 cells. Similarly, GBM10 cells exhibited an increase in G0/G1 and a decrease in S phase under physioxia compared with normoxia. In GBM43 cells, physioxia increased the G0/G1 population while decreasing both S and G2/M phase fractions relative to normoxia. Data are presented as mean ± SEM from three independent experiments (n = 3). P-values calculated by one-way ANOVA where * p< 0.05, ** p<0.01, *** p<0.001, **** p<0.0001

In GBM43 cells, which exhibited an earlier proliferation defect, cell cycle analysis at Day 3 revealed a more pronounced shift in cell cycle distribution under physioxia (Figure 3C). Specifically, GBM43 cells cultured under physioxia displayed an increase in the G0/G1 population, a decrease in S-phase cells, and a reduction in the proportion of cells in G2/M relative to ambient air (Figure 3C). This broader suppression of cycling phases suggests that physioxia induces a more robust cell cycle arrest in GBM43 cells compared with GBM001 and GBM10.

Collectively, these findings indicate that physioxia suppresses glioblastoma cell proliferation by limiting cell cycle progression and reducing S-phase entry, with cell line-specific differences in the extent of cell cycle arrest. Importantly, these cell cycle changes occur in parallel with enhanced migratory capacity under physioxia, suggesting a phenotypic shift away from proliferation and toward motility. Such a proliferation-migration dichotomy has been described in glioblastoma and other invasive cancers and is often linked to transitions toward mesenchymal or EMT-like states.

Given the opposing effects of physioxia on glioblastoma cell proliferation and migration, we next investigated whether physioxia promotes molecular programs associated with epithelial-to-mesenchymal transition and cell motility.

### Figure 4: Physioxia induces Slug expression and correlated with PDGFRβ-associated signaling pathways in a cell line-dependent manner

Previous studies have linked physioxia to the induction of epithelial-to-mesenchymal transition (EMT)-associated transcription factors, including Slug (SNAI2), in glioblastoma and other solid tumors [11]. Increased Slug expression under physioxia has been associated with enhanced migratory and invasive behavior, providing a mechanistic framework for the proliferation-migration dichotomy observed in GBM [11]. Previous studies have identified platelet-derived growth factor receptor β (PDGFRβ) as a key regulator of glioblastoma cell motility through activation of downstream signaling pathways, including AKT and ERK, which have been shown to promote mesenchymal transition and regulate Slug, a critical transcription factor that drives invasive behavior [11]. Although existing evidence suggests that PDGFRβ signaling may primarily regulate Slug through post-transcriptional stabilization of the protein rather than increased transcription, these studies have largely been performed under conventional normoxic culture conditions. Therefore, we sought to determine whether chronic physioxia similarly modulates Slug expression and PDGFRβ pathway activation in patient-derived glioblastoma cells.

**Figure 4:**
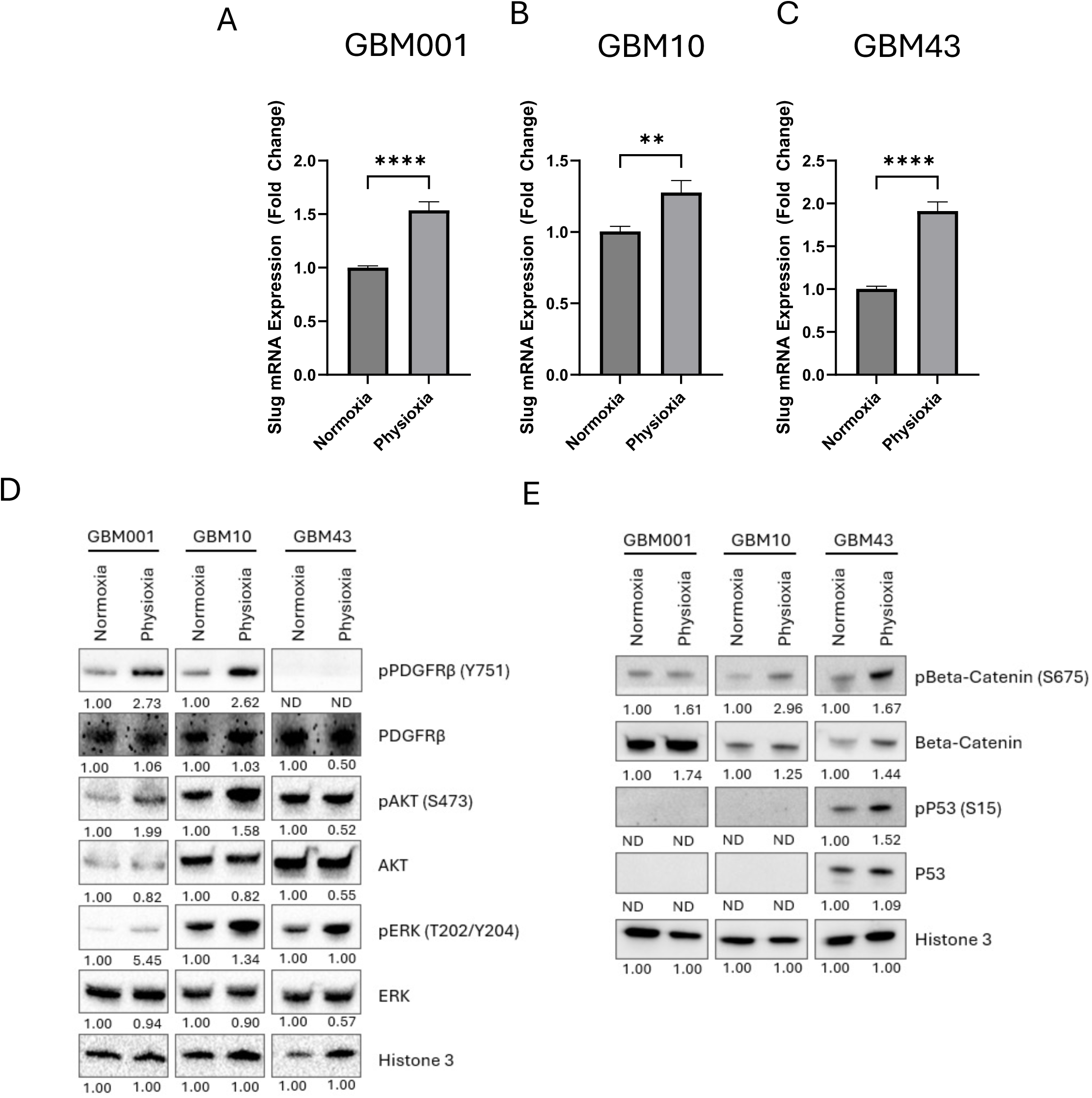
Physioxia increases Slug expression and alters PDGFRβ/AKT/ERK phosphorylation in glioblastoma cells. (A-C) Relative Slug mRNA expression was assessed by RT-qPCR in (A) GBM001, (B) GBM10, and (C) GBM43 cells cultured under normoxic or physioxic conditions. All three cell lines exhibited increased Slug expression in physioxia compared with normoxia. (D) Representative Western blot analysis of GBM001, GBM10, and GBM43 cells cultured under normoxic and physioxic conditions, showing expression of phosphorylated PDGFRβ (Y751), total PDGFRβ, phosphorylated AKT (S473), total AKT, phosphorylated ERK (T202/Y204), and total ERK. Histone H3 was used as a loading control. (E) Representative Western blot analysis of GBM001, GBM10, and GBM43 cells cultured under normoxic and physioxic conditions, showing expression of total P53, pP53 (S15), total β-catenin and pβ-catenin (S675). Histone H3 was used as a loading control. Data represents three independent experiments (n = 3). P-values calculated by one-way ANOVA where ** p<0.01 and **** p<0.0001

RT-qPCR analysis revealed a significant increase in Slug mRNA expression in GBM001, GBM10, and GBM43 cells cultured under physioxia compared with normoxic conditions Figure 4A-C). This upregulation was observed across all three patient-derived glioblastoma lines, indicating that physioxia consistently induces transcriptional programs associated with mesenchymal identity and migration.

To determine whether physioxia-induced Slug expression was associated with upstream signaling pathways, western blot analysis was performed to assess PDGFRβ pathway activity. Protein lysates were probed for total and phosphorylated PDGFRβ (Y751), AKT (S473), and ERK (T202/Y204), with Histone H3 used as a loading control. In both GBM001 and GBM10 cells, physioxia resulted in increased phosphorylation of PDGFRβ at Y751, accompanied by elevated levels of phospho-AKT (S473) and phospho-ERK (T202/Y204), while total protein levels remained largely unchanged. These data indicate that physioxia enhances phosphorylation of PDGFRβ signaling and its downstream AKT and ERK pathways in these cell lines (Figure 4D).

In contrast, GBM43 cells did not exhibit increased changes in PDGFRβ, AKT, or ERK phosphorylation under physioxia compared with normoxia, despite the observed increase in Slug mRNA expression. This lack of differential pathway expression may reflect constitutive signaling activity in GBM43 cells. Notably, GBM43 cells harbor a p53 mutation, which has been reported to promote baseline activation of growth factor signaling pathways, including PI3K/AKT and MAPK/ERK, independent of upstream receptor stimulation. Such constitutive pathway activation may limit further responsiveness to changes in oxygen tension (Figure 4D).

To further investigate the role of p53 signaling under physioxia, western blot analysis was performed for total p53, phospho-p53 (S15), total β-catenin and phospho-β-catenin (S675) in GBM001, GBM10, and GBM43 cells cultured under normoxic and physioxic conditions (Figure 4E). No detectable expression of total p53 or phospho-p53 (S15) was observed in GBM001 or GBM10 cells under either oxygen condition. It is important to note that GBM10 cells do not display a nonsynonymous/missense variant of P53. In contrast, GBM43 cells demonstrated increased phospho-p53 (S15) and phospho-β-catenin (S675) levels under physioxia relative to normoxia. Given that GBM43 harbors a known p53 nonsynonymous/missense mutation, the observed increase in phospho-p53 may reflect stabilization and accumulation of mutant p53 protein rather than activation of canonical wild-type p53 tumor suppressor signaling [21, 28]. Mutant p53 proteins frequently exhibit prolonged protein half-life and aberrant phosphorylation patterns that contribute to oncogenic gain-of-function activities, including enhanced survival, invasion, and adaptation to microenvironmental stress [21]. Increased phosphorylation of β-catenin at S675 under physioxia further suggests activation of pro-survival and pro-migratory signaling pathways in GBM43 cells, as β-catenin signaling has been linked to EMT-associated transcriptional programs and invasive behavior in glioblastoma [29]. Together, these findings support the possibility that physioxia promotes migration in GBM43 cells through mutant p53-associated signaling mechanisms distinct from the PDGFRβ-AKT-ERK axis observed in GBM001 and GBM10 cells.

Together, these findings demonstrate that physioxia induces EMT-associated transcriptional changes across glioblastoma cell lines, while PDGFRβ-AKT-ERK phosphorylation signaling occurs in a cell line-specific manner. In GBM001 and GBM10 cells, enhanced PDGFRβ pathway activity under physioxia provides a potential mechanistic link between increased Slug expression and the pro-migratory phenotype observed under physioxia. In contrast, GBM43 cells may rely on alternative or pre-activated signaling mechanisms to support physioxia-induced migration.

### Figure 5: Physioxia confers cell line-specific resistance to 5-fluorouracil in glioblastoma cells

Physioxia has been reported to influence therapeutic response in glioblastoma and other solid tumors, with multiple studies demonstrating increased resistance to cytotoxic agents under physioxia or hypoxia [11, 14, 25]. Reduced oxygen levels have been shown to promote drug tolerance through mechanisms including altered cell cycle dynamics, enhanced DNA damage response, and activation of pro-survival signaling pathways [30]. These observations provided the rationale to examine whether the physioxia-induced phenotypic changes observed in glioblastoma cells were associated with altered sensitivity to chemotherapy.

**Figure 5:**
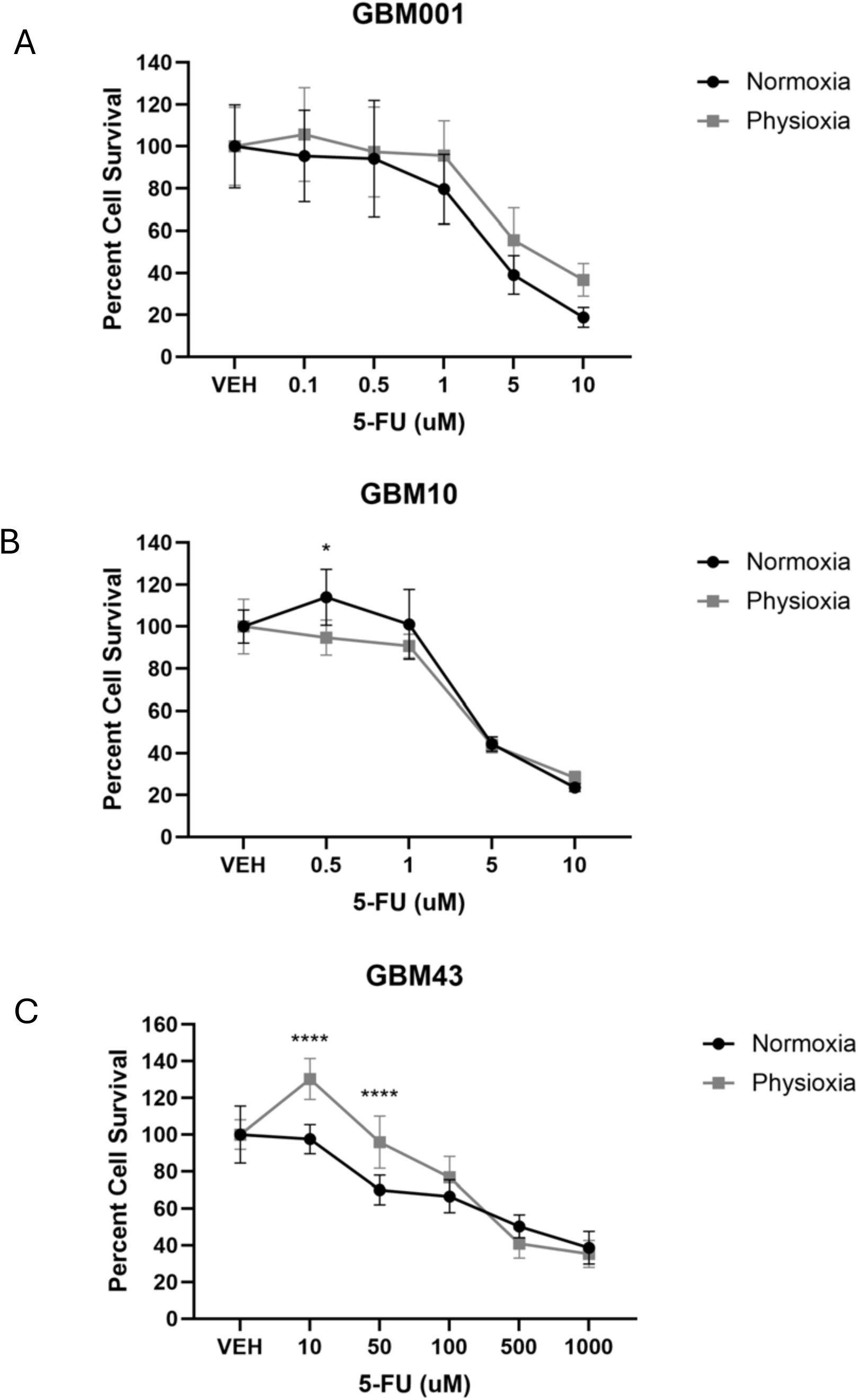
Physioxia differentially modulates chemotherapeutic sensitivity to 5-fluorouracil in glioblastoma cells. (A-C) GBM001 (A), GBM10 (B), and GBM43 (C) cells were treated with increasing concentrations of 5-fluorouracil (5-FU) under normoxic or physioxic conditions. Cell survival was assessed using a BrdU incorporation assay and is expressed as percent survival relative to untreated controls. All cell lines exhibited dose-dependent reductions in survival in response to 5-FU under both oxygen conditions. GBM43 cells displayed increased survival under physioxia at lower 5-FU concentrations, whereas GBM10 cells showed increased survival only at the lowest dose tested. GBM001 cells did not exhibit altered sensitivity to 5-FU under physioxia. Data are presented as mean ± SEM from three independent experiments (n = 3). P-values calculated by one-way ANOVA where * p< 0.05 and **** p<0.0001

To assess the impact of oxygen tension on chemotherapeutic response, GBM001, GBM10, and GBM43 cells were treated with increasing concentrations of 5-fluorouracil (5-FU) under normoxic or physioxic conditions. Cell survival was quantified using a BrdU incorporation assay and expressed as percent survival relative to untreated controls. As expected, all three cell lines exhibited dose-dependent reductions in cell survival in response to 5-FU under both oxygen conditions (Figure 5A-C).

However, distinct differences in chemotherapeutic sensitivity emerged under physioxia in a cell line-dependent manner. GBM001 cells did not demonstrate increased survival under physioxia at any concentration of 5-FU tested, indicating that physioxia does not universally confer chemoresistance across glioblastoma models (Figure 5A). In contrast, GBM10 cells exhibited increased percent cell survival under physioxia only at the lowest tested dose (0.5 µM 5-FU), suggesting a more limited or threshold-dependent resistance phenotype (Figure 5B). GBM43 cells cultured under physioxia displayed a significant increase in percent cell survival compared with normoxia at 10 µM and 50 µM 5-FU, indicating reduced sensitivity to higher drug concentrations (Figure 5C).

These data demonstrate that physioxia can promote resistance to 5-FU in glioblastoma cells, but that this effect is highly dependent on cellular context. The observed resistance patterns parallel earlier findings of physioxia-induced cell cycle arrest and pathway activation, suggesting that reduced proliferative activity and enhanced survival signaling may contribute to decreased chemotherapy sensitivity in select glioblastoma lines.

Together with the enhanced migratory capacity and altered cell cycle dynamics observed under physioxia, these findings support a model in which physioxia promotes a more invasive and therapy-resistant phenotype in a subset of glioblastoma cells. The cell line-specific nature of this response highlights the importance of genetic and signaling context in determining therapeutic vulnerability within the physioxic tumor microenvironment.

### Figure 6: Glioblastoma cells maintained under continuous physioxia exhibit enhanced proliferation and migration

Previous studies have demonstrated that exposure of primary tumor cells to ambient air can induce rapid and potentially irreversible biological changes [7, 8, 25]. Notably, work by Broxmeyer and colleagues has shown that brief exposure to normoxic conditions, in a matter of minutes, can alter transcriptional programs, signaling pathways, and functional behavior of cells that normally reside in low-oxygen niches [8, 25]. These findings underscore the importance of maintaining physioxia during tumor cell isolation and culture to preserve native cellular phenotypes.

**Figure 6:**
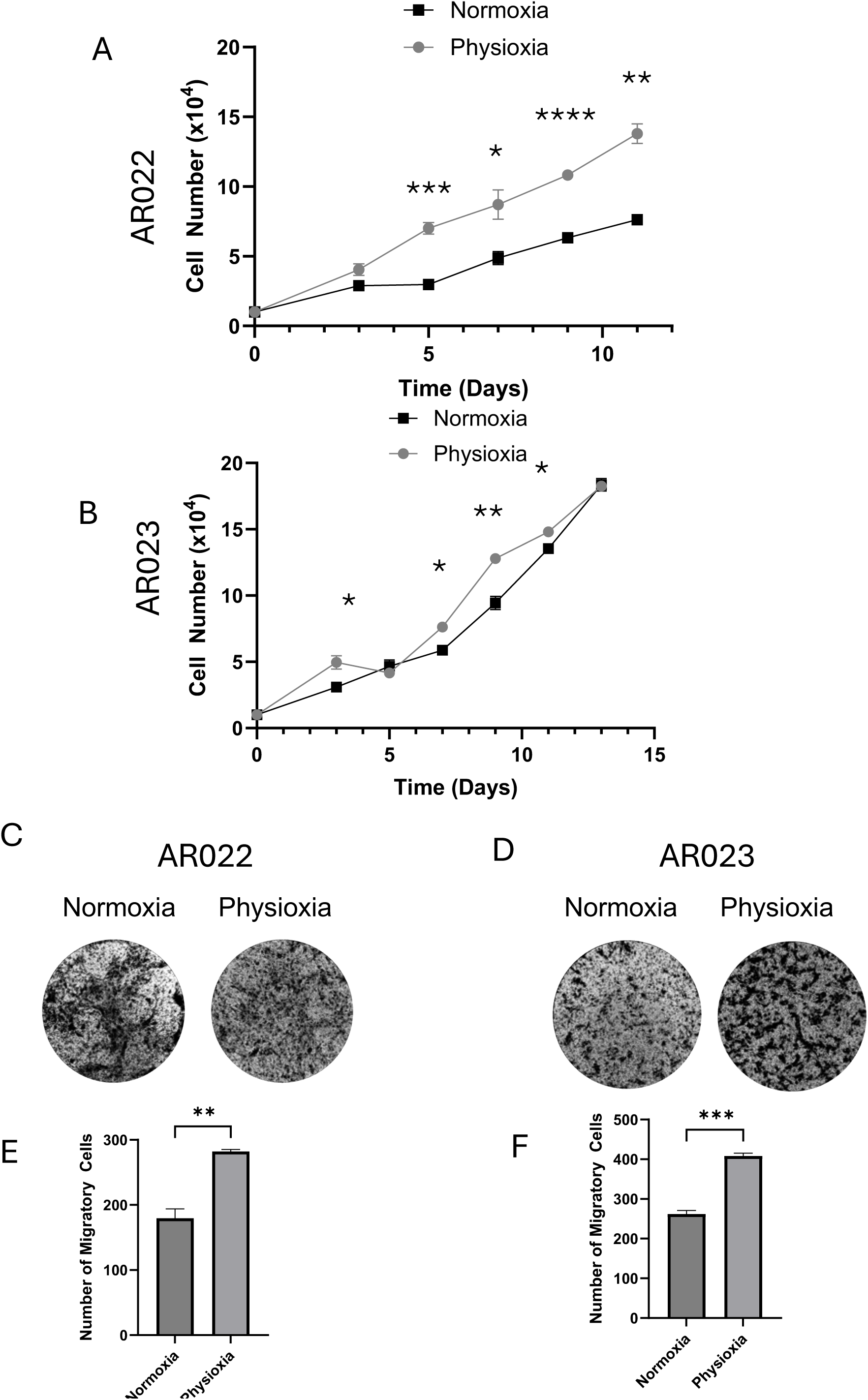
Patient-derived glioblastoma cultures retain enhanced proliferative and migratory phenotypes under physioxia. (A-B) Cell proliferation of patient-derived glioblastoma cultures (A) AR022 and (B) AR023 was assessed over a 14-day period following establishment under physioxic conditions and subsequent culture in either normoxia or physioxia. Both AR022 and AR023 cells exhibited increased proliferation under physioxia compared with normoxia. (C-D) Representative images of Transwell migration assays for (C) AR022 and (D) AR023 cells cultured under normoxic or physioxic conditions. (E-F) Quantitative analysis of migrated cells demonstrating significantly increased migration under physioxia in (E) AR022 and (F) AR023 cultures. Data are presented as mean ± SEM from three independent experiments (n = 3). P-values calculated by one-way ANOVA where * p< 0.05, ** p<0.01, *** p<0.001, **** p<0.0001

In the present study, we compared two distinct culture paradigms to assess the impact of continuous physioxia on glioblastoma cell behavior. Patient-derived tumor samples AR022 and AR023 were collected, processed, and maintained entirely under physioxic conditions and compared with parallel cultures acclimated to normoxia. In contrast, the GBM001, GBM10, and GBM43 cells used in Figures 1-5 were originally derived from the physiological oxygen environment of the brain but were subsequently cultured long-term under ambient air and later re-exposed to physioxia. This distinction is critical given prior reports that even transient normoxic exposure may permanently reprogram tumor cell behavior.

Analysis of patient-derived AR022 and AR023 cells revealed a significant increase in cell proliferation when maintained under physioxia compared with normoxia-acclimated cultures (Figure 6A-B). In addition to enhanced growth, both AR022 and AR023 cells exhibited increased migratory capacity under physioxic conditions, as assessed by transwell migration assays (Figure 6C-F). Notably, the physioxia-induced increase in migration observed in these continuously physioxia-maintained patient-derived cells mirrored the enhanced migratory phenotype seen in GBM001, GBM10, and GBM43 cells under physioxia (Figures 1-5), despite differences in culture history.

These findings demonstrate that physioxia promotes a pro-tumorigenic phenotype in glioblastoma cells regardless of whether cells are continuously maintained under physioxia or reintroduced to physiological oxygen levels after normoxic culture. Importantly, the observation that AR022 and AR023 cells display increased proliferation under physioxia contrasts with the reduced proliferation observed in GBM001, GBM10, and GBM43 cells, suggesting that long-term normoxic adaptation may alter the proliferative response to physioxia.

Together, these results highlight the importance of modeling physiological oxygen tension throughout tumor cell isolation and culture. The enhanced proliferation and migration observed in AR022 and AR023 cells maintained under continuous physioxia suggest that traditional normoxic culture conditions may obscure or distort native glioblastoma phenotypes. By demonstrating that physioxia consistently enhances migratory behavior across multiple glioblastoma models while differentially regulating proliferation depending on culture history, this work emphasizes the need for physiologically relevant oxygen conditions during biospecimen processing and downstreatm applications to accurately study glioblastoma biology and therapeutic response.

## Discussion

Glioblastoma develops within a physiologically low-oxygen microenvironment that is not accurately reproduced by standard culture conditions. In this study, we show that prolonged culture under physioxia (5% O□) alters multiple aspects of glioblastoma cell behavior, including migration, proliferation, cell cycle progression, signaling pathway activation, EMT-associated transcription, and therapeutic response. Collectively, these findings support a model in which physiological oxygen tension promotes a more invasive and adaptive glioblastoma phenotype while differentially regulating proliferation and drug sensitivity depending on cellular context.

A consistent finding across all glioblastoma models examined was enhanced migration under physioxia. GBM001, GBM10, and GBM43 cells all exhibited significantly increased transwell migration despite differences in genetic background, signaling responses, and proliferative behavior. These observations extend previous reports from other tumor systems to patient-derived glioblastoma models and indicate that increased motility is a conserved response to physiological oxygen tension. Given the highly infiltrative nature of GBM in vivo, these findings suggest that physioxia favors phenotypes associated with invasion and tissue adaptation rather than maximal proliferative output.

In contrast to the increase in migration, physioxia reduced long-term proliferation in GBM001, GBM10, and GBM43 cells. Reduced cell accumulation was associated with increased G0/G1 occupancy and reduced S-phase entry, indicating suppression of cell cycle progression under physiological oxygen conditions. GBM43 cells exhibited the strongest response, including reductions in both S-phase and G2/M populations, consistent with a more pronounced cell cycle arrest. Together, these data support a proliferation-migration dichotomy in glioblastoma under physioxia, in which invasive behavior is enhanced while proliferative activity is restrained [31]. Such a shift may provide a selective advantage within the brain microenvironment, where adaptation to metabolic stress and therapeutic pressure is required for tumor persistence.

At the molecular level, physioxia increased Slug (SNAI2) expression in all glioblastoma lines examined, linking physiological oxygen tension to EMT-associated transcriptional programs. Slug has been implicated in mesenchymal transition, cytoskeletal remodeling, and invasion in glioblastoma and other cancers [11]. The consistent induction of Slug across all models suggests that activation of mesenchymal transcriptional programs represents a conserved response to physioxia. In addition to Slug, hypoxia-responsive pathways have been shown to promote glioblastoma migration through upregulation of ODZ1 (TENM1), a transmembrane protein that enhances actin cytoskeletal remodeling and cell migration downstream of hypoxia-inducible signaling [32]. Together, these findings suggest that activation of mesenchymal and pro-migratory transcriptional programs represents a conserved cellular response to reduced oxygen availability.

Analysis of upstream signaling pathways demonstrated substantial cell line-dependent differences. In GBM001 and GBM10 cells, physioxia increased phosphorylation of PDGFRβ, AKT, and ERK, consistent with activation of signaling pathways associated with glioblastoma migration and mesenchymal transition [11]. These findings support a mechanistic relationship between physiological oxygen tension and PDGFRβ-dependent signaling in promoting invasive behavior.

In contrast, GBM43 cells showed increased Slug expression and migration without detectable increases in PDGFRβ, AKT, or ERK phosphorylation. One explanation for this divergence is the presence of mutant p53 in GBM43 cells. Mutant p53 has been reported to promote constitutive PI3K/AKT and MAPK/ERK signaling and to uncouple EMT-associated transcriptional programs from receptor tyrosine kinase activation [21, 29]. Baseline pathway activation may therefore limit further inducibility under physioxia. Consistent with this interpretation, physioxia increased phospho-p53 (S15) and phospho-β-catenin (S675) levels in GBM43 cells, suggesting engagement of alternative signaling mechanisms linked to invasion and cellular adaptation. Increased β-catenin signaling has previously been associated with EMT-related transcription and invasive behavior in glioblastoma [29].

The pronounced cell cycle arrest observed in GBM43 cells may also contribute to the distinct signaling phenotype observed under physioxia. Quiescent glioblastoma cells can retain invasive capacity while displaying reduced dependence on growth factor-mediated proliferative signaling [33]. This may explain why GBM43 cells maintained enhanced migration without additional PDGFRβ-associated pathways. Together, these findings indicate that although physioxia consistently promotes migration, the signaling pathways supporting this phenotype are strongly influenced by underlying genetic context, including p53 status.

Physioxia also altered therapeutic response in a cell line-dependent manner. GBM43 cells exhibited increased resistance to 5-fluorouracil (5-FU) at higher drug concentrations, while GBM10 cells showed limited resistance only at low-dose treatment. In contrast, GBM001 cells did not exhibit increased survival under physioxia. The observed resistance patterns correlated with reduced proliferation and altered cell cycle progression, consistent with studies linking quiescence and reduced S-phase entry to decreased sensitivity to antimetabolite therapies. These findings suggest that physiological oxygen tension can promote a therapy-tolerant state in specific glioblastoma contexts but does not uniformly induce chemoresistance.

An important observation from this study is the influence of culture history on glioblastoma behavior under physioxia. GBM001, GBM10, and GBM43 cells were initially derived from the low-oxygen brain environment but subsequently maintained long-term under ambient air before re-exposure to physioxia. Previous studies by Broxmeyer and colleagues demonstrated that even brief exposure to atmospheric oxygen can induce rapid and potentially irreversible cellular alterations [8, 25]. To address this issue, we examined patient-derived AR022 and AR023 cells that were isolated and continuously maintained under physioxia. In contrast to normoxia-adapted GBM lines, these cells displayed increased proliferation and migration under physioxic conditions. While enhanced migration was consistent across all models, the increased proliferation observed in continuously physioxic cultures contrasted with the growth suppression seen in normoxia-adapted lines. These findings suggest that prolonged exposure to ambient air may alter glioblastoma responses to physiological oxygen tension, particularly with respect to proliferative behavior.

Overall, our findings demonstrate that physiological oxygen tension promotes an invasive and adaptive glioblastoma phenotype characterized by enhanced migration, EMT-associated transcription, altered cell cycle progression, and context-dependent therapeutic resistance. These results further indicate that oxygen tension is a major determinant of glioblastoma cell state and that conventional normoxic culture conditions may obscure biologically relevant phenotypes. The molecular signature identified in this study differs from that reported in previous investigations of hypoxia-induced glioblastoma invasion, which have implicated additional mediators such as ODZ1 (TENM1), a HIF-responsive regulator of actin cytoskeletal remodeling and cell migration, as well as other canonical hypoxia-induced targets associated with severe oxygen deprivation. These differences may reflect the distinct oxygen paradigm employed in our study. Whereas many previous investigations have examined acute, severe hypoxia (typically ≤1-2% O for 24-72 hours), we modeled chronic physiological oxygen tension (5% O□), which more closely approximates the oxygen environment experienced by normal brain tissue. Short-term exposure to reduced oxygen may not allow tumor cells to fully establish long-term adaptive programs, while prolonged culture under atmospheric oxygen before hypoxic exposure may induce persistent molecular alterations that influence subsequent responses to reduced oxygen tension.

Collectively, our findings highlight the importance of oxygen history as an often-overlooked experimental variable and suggest that oxygen tension during tissue acquisition, cell isolation, expansion, and experimentation can substantially influence the molecular and phenotypic readouts commonly used in cancer biology. Further studies are therefore needed to define how both oxygen concentration and duration of exposure shape glioblastoma biology and to establish physiologically relevant culture conditions that more accurately model the native tumor microenvironment. By modeling glioblastoma under sustained physiological oxygen tension, experimental systems may better recapitulate tumor biology, improve the reproducibility and translational relevance of preclinical studies, and ultimately provide a more accurate platform for evaluating therapeutic responses.

### Future directions

Several important questions emerge from this work. First, future studies should define the precise molecular mechanisms linking physioxia to Slug induction, including the relative contributions of hypoxia-inducible factors, chromatin remodeling, and non-canonical signaling pathways. Second, systematic comparisons of continuously physioxia-maintained versus normoxia-adapted glioblastoma cultures will be essential to determine which phenotypes represent true in vivo behavior. Third, given the cell line-specific signaling responses observed here, integrating physioxia modeling with genomic and epigenomic profiling may reveal biomarkers predictive of invasion or therapy resistance. Finally, evaluating standard-of-care therapies, including temozolomide and radiation, under physiological oxygen conditions will be critical for improving the translational relevance of preclinical GBM studies.

In summary, this study demonstrates that physioxia is a central regulator of glioblastoma cell behavior, shaping migration, proliferation, signaling, and therapeutic response. Incorporating physioxia into experimental design will be essential for accurately modeling glioblastoma biology and for developing more effective therapeutic strategies.

## Conclusions

This study demonstrates that sustained culture under physiological oxygen tension fundamentally alters glioblastoma cell behavior, promoting enhanced migration, altered cell cycle progression, differential activation of signaling pathways, and context-dependent therapeutic sensitivity. While physioxia consistently increased migratory capacity across all glioblastoma models examined, its effects on proliferation depended on the cells’ oxygen history, highlighting the importance of prior culture conditions in shaping tumor phenotypes. These findings establish oxygen tension as an active regulator of glioblastoma biology rather than simply a culture parameter and suggest that conventional atmospheric oxygen conditions may obscure clinically relevant cellular behaviors.

Collectively, our results support the incorporation of physiologically relevant oxygen conditions throughout glioblastoma cell isolation, culture, and experimentation to improve the biological accuracy and translational value of preclinical models. Accounting for both oxygen concentration and oxygen history may enhance the reproducibility of experimental findings, provide more faithful models of the tumor microenvironment, and improve the evaluation of therapeutic strategies for glioblastoma.

---

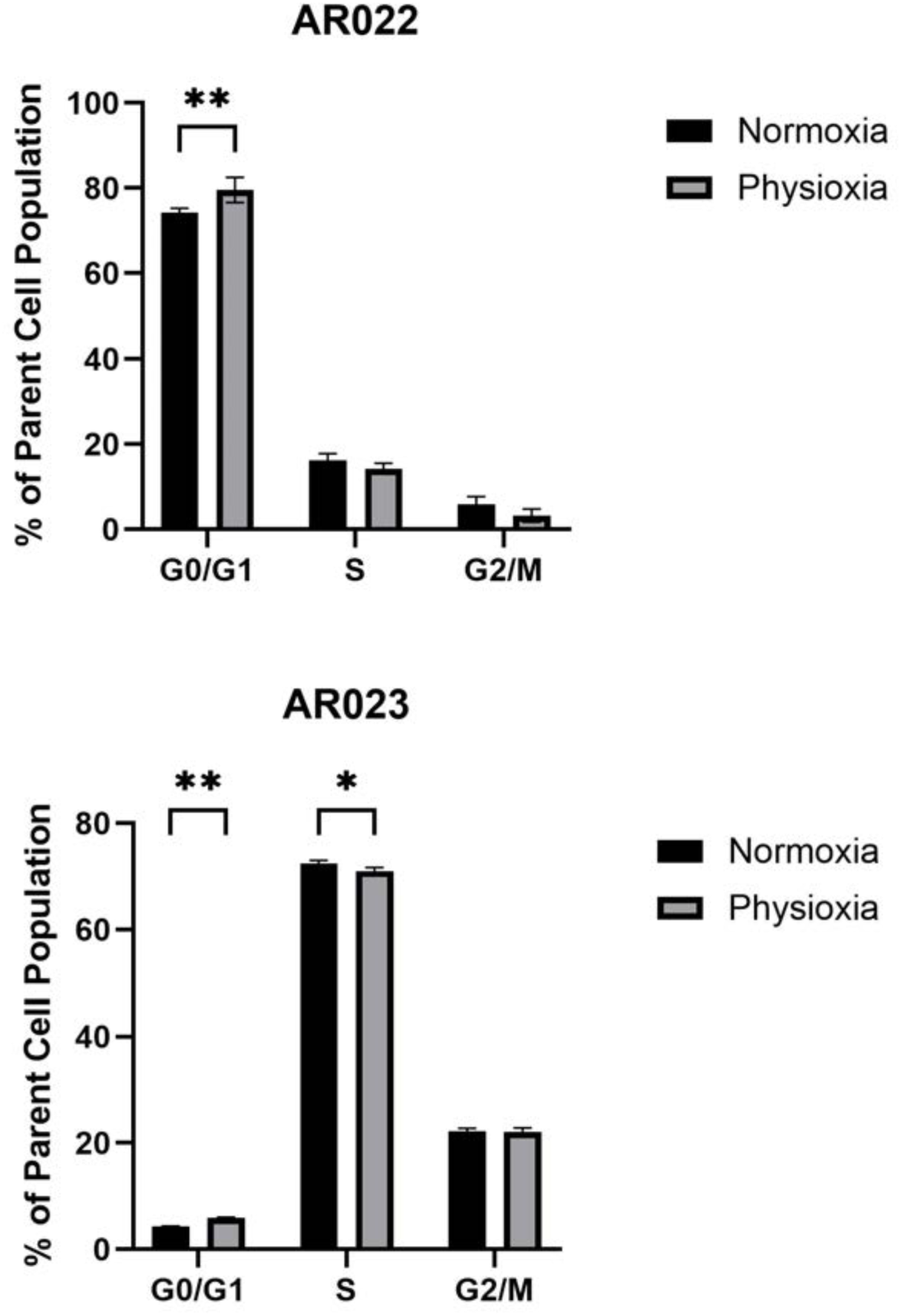

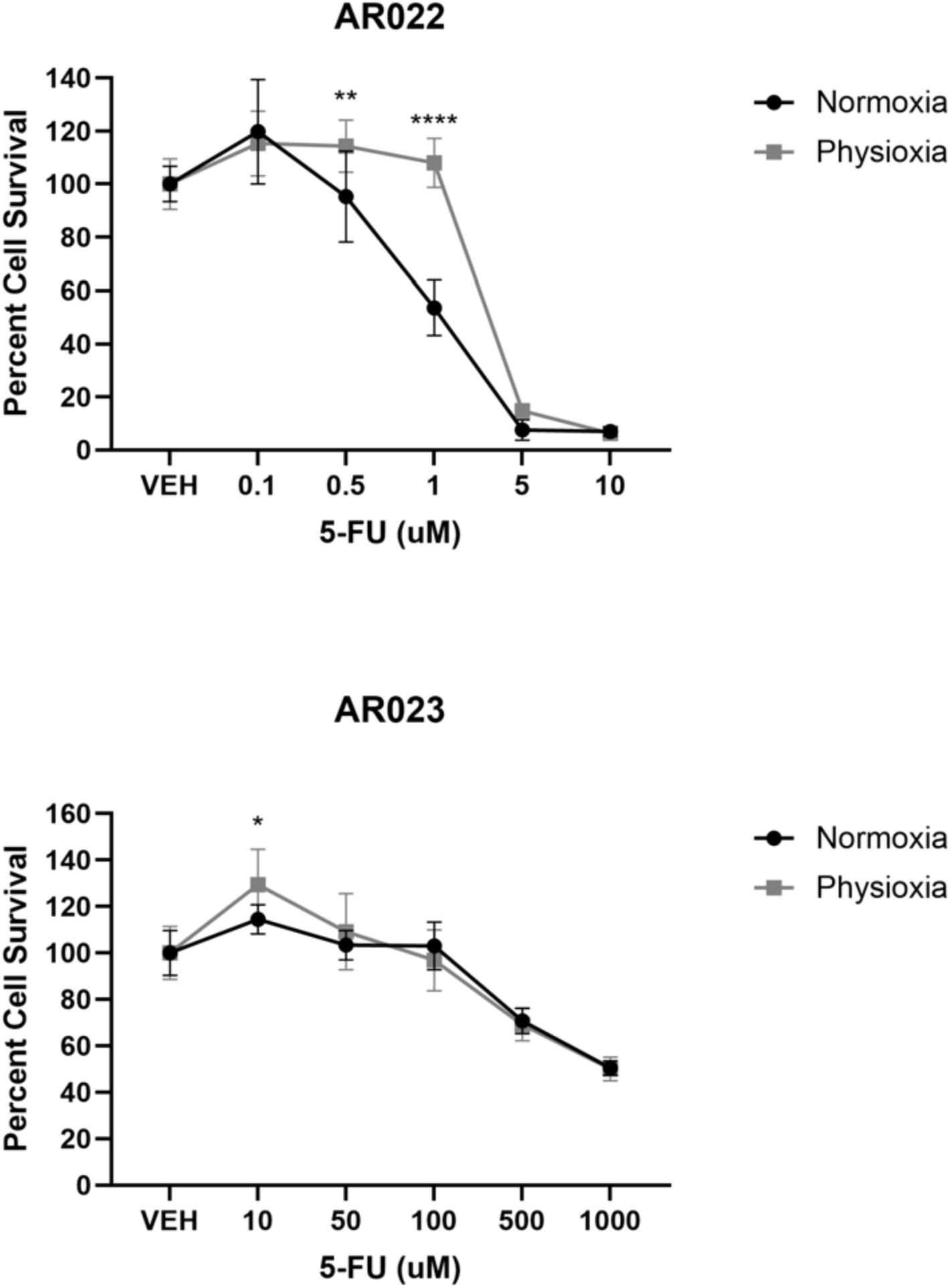

## Notes

### Competing Interest Statement

The authors have declared no competing interest.

## References

1. Sipos, D., et al., Glioblastoma: Clinical Presentation, Multidisciplinary Management, and Long-Term Outcomes. Cancers (Basel), 2025. 17(1).

2. Wu, W., et al., Glioblastoma multiforme (GBM): An overview of current therapies and mechanisms of resistance. Pharmacol Res, 2021. 171: p. 105780.

3. Michelucci, A., et al., Hypoxia, Ion Channels and Glioblastoma Malignancy. Biomolecules, 2023. 13(12).

4. Park, J.H. and H.K. Lee, Current Understanding of Hypoxia in Glioblastoma Multiforme and Its Response to Immunotherapy. Cancers (Basel), 2022. 14(5).

5. Alva, R., et al., Supraphysiological Oxygen Levels in Mammalian Cell Culture: Current State and Future Perspectives. Cells, 2022. 11(19).

6. Jagannathan, L., S. Cuddapah, and M. Costa, Oxidative stress under ambient and physiological oxygen tension in tissue culture. Curr Pharmacol Rep, 2016. 2(2): p. 64–72.

7. Broxmeyer, H.E., et al., The importance of hypoxia and extra physiologic oxygen shock/stress for collection and processing of stem and progenitor cells to understand true physiology/pathology of these cells ex vivo. Curr Opin Hematol, 2015. 22(4): p. 273–8.

8. Mantel, C.R., et al., Enhancing Hematopoietic Stem Cell Transplantation Efficacy by Mitigating Oxygen Shock. Cell, 2015. 161(7): p. 1553–65.

9. Druker, J., et al., Role of Hypoxia in the Control of the Cell Cycle. Int J Mol Sci, 2021. 22(9).

10. Garcia, D., et al., Short exposure to hyperoxia causes cultured lung epithelial cell mitochondrial dysregulation and alveolar simplification in mice. Pediatr Res, 2021. 90(1): p. 58–65.

11. Adebayo, A.K., et al., Oxygen tension-dependent variability in the cancer cell kinome impacts signaling pathways and response to targeted therapies. iScience, 2024. 27(6): p. 110068.

12. Sun, X., et al., Physiologically normal 5% O2 supports neuronal differentiation and resistance to inflammatory injury in neural stem cell cultures. J Neurosci Res, 2015. 93(11): p. 1703–12.

13. Warpsinski, G., et al., Nrf2-regulated redox signaling in brain endothelial cells adapted to physiological oxygen levels: Consequences for sulforaphane mediated protection against hypoxia-reoxygenation. Redox Biol, 2020. 37: p. 101708.

14. Adebayo, A.K. and H. Nakshatri, Modeling Preclinical Cancer Studies under Physioxia to Enhance Clinical Translation. Cancer Res, 2022. 82(23): p. 4313–4321.

15. Liang, Y., J. Liu, and Z. Feng, The regulation of cellular metabolism by tumor suppressor p53. Cell Biosci, 2013. 3(1): p. 9.

16. Riley, T., et al., Transcriptional control of human p53-regulated genes. Nat Rev Mol Cell Biol, 2008. 9(5): p. 402–12.

17. Zhang, Y., et al., The p53 Pathway in Glioblastoma. Cancers (Basel), 2018. 10(9).

18. Zhang, C., et al., The Interplay Between Tumor Suppressor p53 and Hypoxia Signaling Pathways in Cancer. Front Cell Dev Biol, 2021. 9: p. 648808.

19. Kim, Y., et al., Platelet-derived growth factor receptors differentially inform intertumoral and intratumoral heterogeneity. Genes Dev, 2012. 26(11): p. 1247–62.

20. Li, Q.L., et al., Activation of PI3K/AKT and MAPK pathway through a PDGFRbeta-dependent feedback loop is involved in rapamycin resistance in hepatocellular carcinoma. PLoS One, 2012. 7(3): p. e33379.

21. Weissmueller, S., et al., Mutant p53 drives pancreatic cancer metastasis through cell-autonomous PDGF receptor beta signaling. Cell, 2014. 157(2): p. 382–394.

22. Yun, Z. and P.M. Glazer, Tumor suppressor p53 stole the AKT in hypoxia. J Clin Invest, 2015. 125(6): p. 2264–6.

23. Hockaden, N., et al., Amyloidogenesis promotes HSF1 activity enhancing cell survival during breast cancer metastatic colonization. Cell Stress Chaperones, 2025. 30(3): p. 143–159.

24. Downing, N.F., K.M. Mills, and P.C. Hollenhorst, PRC2/FOXO1-Mediated Repression Determines Interchangeability of ETS Oncogenes in Prostate Cancer and Ewing Sarcoma. Mol Cancer Res, 2026. 24(1): p. 48–59.

25. Kumar, B., et al., Tumor collection/processing under physioxia uncovers highly relevant signaling networks and drug sensitivity. Sci Adv, 2022. 8(2): p. eabh3375.

26. Semenza, G.L., Molecular mechanisms mediating metastasis of hypoxic breast cancer cells. Trends Mol Med, 2012. 18(9): p. 534–43.

27. Obeagu, E.I., Oxygen gradients in tumor tissues implications for breast cancer metastasis - a narrative review. Ann Med Surg (Lond), 2025. 87(6): p. 3372–3380.

28. Rivlin, N., et al., Mutations in the p53 Tumor Suppressor Gene: Important Milestones at the Various Steps of Tumorigenesis. Genes Cancer, 2011. 2(4): p. 466–74.

29. Alvarado-Ortiz, E., et al., Mutant p53 Gain-of-Function: Role in Cancer Development, Progression, and Therapeutic Approaches. Front Cell Dev Biol, 2020. 8: p. 607670.

30. Devi, D., et al., Hypoxia-driven metabolic and molecular reprogramming: From tumor microenvironment to therapeutic interventions. Pathol Res Pract, 2025. 276: p. 156271.

31. Hatzikirou, H., et al., ’Go or grow’: the key to the emergence of invasion in tumour progression? Math Med Biol, 2012. 29(1): p. 49–65.

32. Velasquez, C., et al., Hypoxia Can Induce Migration of Glioblastoma Cells Through a Methylation-Dependent Control of ODZ1 Gene Expression. Front Oncol, 2019. 9: p. 1036.

33. Xie, X.P., et al., Quiescent human glioblastoma cancer stem cells drive tumor initiation, expansion, and recurrence following chemotherapy. Dev Cell, 2022. 57(1): p. 32–46 e8.

